# Brain injury degrades the precision of hippocampal spatial representations

**DOI:** 10.64898/2026.08.28.747393

**Authors:** Alexa Tierno, Robert F Hunt

## Abstract

Traumatic brain injury (TBI) disrupts spatial memory, but the circuit mechanisms are unknown. Here we used *in vivo* endoscopic calcium imaging to record CA1 activity dynamics in freely behaving mice with TBI. Spontaneous CA1 neuron activity is broadly preserved three months after brain injury. However, chronic TBI reorganizes hippocampal spatial representations by increasing the fraction of spatially unresponsive neurons, broadening place fields among spatially tuned neurons and diminishing the persistence of spatial representations across sessions. These circuit alterations are accompanied by impaired discrimination of a novel spatial environment. Thus, TBI reorganizes the cellular structure and precision of hippocampal spatial representations despite preserved overall CA1 activity, revealing a dissociation between neuronal activity and network function that may underlie persistent cognitive dysfunction.

## Introduction

The hippocampus generates dynamic representations of space that are essential for navigation and memory. In CA1, pyramidal neurons fire selectively at specific locations within an environment, collectively forming a neural representation of space whose precision and stability depend on coordinated interactions across local and long-range circuits (O’Keefe and Dostrovsky, 1971; O’Keefe, 1976; Bittner et al., 2015). Entorhinal inputs provide major excitatory drive into hippocampus, while local inhibitory circuits regulate network excitability and constrain the spatial tuning of CA1 neurons (Klausberger et al., 2003; Solstad et al., 2006; Brun et al., 2008; Geiller et al., 2022; Zutshi et al., 2022). Disruption of these circuits can alter the precision, information content or stability of spatial representations in hippocampus (Morris et al., 1982; Calton et al., 2003; Hales et al., 2014; Bittner et al., 2015; Udakis et al., 2025).

Traumatic brain injury (TBI) produces extensive and persistent remodeling of hippocampal circuitry, including loss of inhibition, fewer entorhinal inputs and increased local excitatory connectivity (Toth et al., 1997; Santhakumar et al., 2000; Hunt et al., 2009, 2011; Frankowski et al., 2022). Electrophysiological studies have provided preliminary evidence of altered task-related hippocampal coding after TBI (Eakin and Miller, 2012; Broussard et al., 2020), but how hippocampal spatial representations are organized once chronic circuit remodeling has emerged is unknown. Work in other neurodegenerative disorders affecting hippocampus, such as epilepsy and Alzheimer’s disease models, has applied more classic paradigms and shown that circuit remodeling can degrade spatial coding (Liu et al., 2003; Cacucci et al., 2008; Mably et al., 2017; Jun et al., 2020; Shuman et al., 2020). This raises the question of whether chronic TBI reshapes how the hippocampus represents space.

Here, we used longitudinal endoscopic calcium imaging to examine CA1 neuronal activity and spatial representations in freely behaving mice three months after TBI. We first characterized spontaneous neuronal activity, then assessed the recruitment, spatial precision and persistence of CA1 place cell representations during a linear track task. Finally, we tested the behavioral consequences of these circuit changes using a spatial novelty discrimination task. We find that chronic TBI leaves overall CA1 neuronal activity broadly preserved but disrupts its organization into spatial representations, with reduced place cell recruitment, broader place fields and diminished persistence across sessions. These circuit changes are accompanied by impaired discrimination of novel spatial environments. Together, these results identify degradation of hippocampal network precision as a circuit correlate of cognitive dysfunction after TBI.

## Methods

### Experimental model

All animal procedures were approved by the Institutional Animal Care and Use Committee of University of California Irvine and performed in accordance with National Institutes of Health Guidelines for the Care and Use of Laboratory Animals. Experiments were performed in adult homozygous *Camk2a-Cre* mice (JAX Stock No. 005359) of both sexes maintained under standard housing conditions on a 12 h light/dark cycle with food and water available *ad libitum* until food restriction was initiated for linear track training.

### Brain injury

Controlled cortical impact (CCI) was performed in male and female mice at postnatal day 60–65 as previously described (Frankowski et al., 2022). Briefly, mice were anesthetized with 2% isoflurane and secured in a custom stereotaxic frame. Following a midline scalp incision, a 4–5 mm craniotomy was made approximately 1 mm lateral to the sagittal suture and centered between bregma and lambda. The skull cap was removed with fine forceps without disrupting the underlying dura. CCI was delivered to the neocortex using a computer-controlled, pneumatically driven impactor fitted with a 3 mm diameter beveled stainless-steel tip (Precision Systems and Instrumentation, TBI-0310) at a depth of 1.0 mm, velocity of 3.5 m/s, and dwell time of 500 ms. The incision was sutured without replacing the skull cap, which was retained for subsequent lens implantation. Animals recovered on a heating pad. Buprenorphine hydrochloride (Buprenex; 0.05 mg/kg, intraperitoneally) was administered at the time of surgery and 12 h later.

### Virus injection

AAV1-syn-FLEX-jGCAMP8m-WPRE (Addgene, plasmid no. 162378; 1.0 × 10^13 genome copies/mL) was diluted 1:10 in sterile PBS immediately before injection. Virus was loaded into beveled glass micropipettes (50 μm tip diameter; Wiretrol 5 μm, Drummond Scientific), and 100 nL was injected at ∼15 nL/min. The micropipette was left in place for 10 min before withdrawal. Viral injections targeting dorsal CA1 were performed 5–8 weeks after CCI. Injection coordinates were established in preliminary experiments to optimize targeting of dorsal CA1 and to determine viral titer and injection volume for pyramidal cell specificity (**Fig. S1**). Coordinates for control mice were anteroposterior (AP) −2.1 mm, mediolateral (ML) 1.5 mm, and dorsoventral (DV) −1.2 mm; coordinates for CCI mice were AP −1.9 mm, ML 1.5 mm, and DV −0.65 mm.

### GRIN lens implantation

Three weeks after viral injection, a ∼1 mm craniotomy was made above the injection site. In CCI mice, the skull cap was replaced at this time. Cortical tissue was carefully aspirated using 27-and 30-gauge blunt needles connected to a vacuum pump until the alveus overlying CA1 was visible. Cold sterile saline was continuously applied during aspiration. Surgifoam (Ethicon) was used to control bleeding and establish a clear optical window. A GRIN lens (1 mm diameter, 4 mm length; Inscopix, 1050-004595) was slowly lowered to the target region using a stereotaxic arm (DV −1.2 mm for control mice; −0.85 mm for brain injured mice to account for hippocampal herniation). The lens was secured to the skull using cyanoacrylate glue and dental cement. Kwik-Sil (World Precision Instruments) was applied to the exposed lens surface to protect the optical interface. One week later, mice were briefly anesthetized with isoflurane and a miniscope mounted on a baseplate was positioned over the GRIN lens. The miniscope was adjusted to identify an optimal field of view (FOV) containing in-focus jGCaMP8m-expressing cells. The baseplate was secured using cyanoacrylate glue and dental cement. The miniscope was removed and replaced with a protective plastic cap between recording sessions.

### Linear track

One week after lens implantation, mice were food restricted to approximately 85% of their initial body weight. Body weight and general health were monitored daily, and food was provided if signs of fatigue or distress were observed. Mice were handled for at least 5 min/day for 2 weeks before behavioral testing. To habituate mice to the weight of the miniscope, a dummy scope (OpenEphys, OEPS-7409) was worn for progressively increasing periods beginning 1 week before experiments, until mice could comfortably tolerate the scope for 15 min/day. The linear track consisted of a 200 × 6 × 12 cm white acrylic track with black visual cues on the walls (**Fig. S2**). Room illumination was maintained at 20 lux. Before each recording session, the miniscope was attached to the baseplate and the FOV was confirmed. Home-cage recordings were obtained on the first day of linear track training immediately before the first track session. Mice were placed at one end of the track and trained to run back and forth for a food reward (sugar pellet; Bio-Serv, F0023) after completion of each lap. Each imaging session consisted of at least 15 laps, with four sessions per day separated by 10 min home-cage rest periods. Training was conducted for 7 days, and data from day 7 were used for spatial analyses. The track was cleaned with 70% ethanol between trials.

### Calcium imaging

Calcium imaging was performed using UCLA Miniscope v4 systems (http://miniscope.org) connected to a data acquisition system through a flexible coaxial cable (OpenEphys, OEPS-5503). The acquisition system was connected to a computer through USB 3.0 (**Fig. S3**). The focal plane and excitation intensity were adjusted before each recording session. Miniscope acquisition was controlled using a Bonsai workflow (Lopes et al., 2015), and images were acquired at 20 frames/s.

### Position and speed tracking

Behavior was recorded using an overhead camera (acA2040-90um, Basler) at 30 frames/s and synchronized to miniscope imaging frames using hardware timestamps in a custom Bonsai workflow (**Fig. S3**). The animal’s position was tracked offline using the miniscope-mounted LED and DeepLabCut (Mathis et al., 2018). Tracking output was visually inspected, and erroneous coordinates were corrected manually. Position data were smoothed using a Gaussian filter (σ = 150 ms), and running speed was calculated from the smoothed position data.’

### Calcium imaging pre-processing

Calcium imaging data were processed using MIN1PIPE for background subtraction, motion correction, calcium trace extraction, and spatial footprint extraction (Lu et al., 2018). Calcium events were inferred from fluorescence traces using MLspike (Deneux et al., 2016) and converted to firing rates. Traces with high noise levels (σ > 0.1; σ defined as the root-mean-square of fluorescence signals between 3 and 20 Hz) were excluded. Mice in which >30% of extracted cells were excluded were removed from subsequent analyses (**Fig. S4**). Spatial footprints generated by MIN1PIPE were used to register cells across sessions using CellReg (Sheintuch et al., 2017). Automated cell extraction, spike inference, and cell registration were manually inspected for accuracy.

### Place cell analysis

For linear track analyses, only periods during which mice were running >5 cm/s were included. Rightward and leftward traversals were analyzed separately. Position was divided into 2 cm spatial bins, and firing rate was calculated for each bin as the number of inferred calcium events divided by occupancy time. Rate maps were smoothed using a Gaussian kernel (σ = 2 cm). Spatial information was calculated for each neuron using the method of Skaggs et al. (1993):

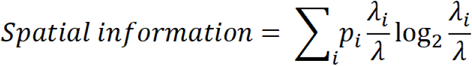

where *λ_i_* is the mean firing rate in spatial bin *i*, *λ* is the overall mean firing rate, and *p_i_* is the probability of occupancy of spatial bin *i*.

Place cells were identified using a circular-shuffling procedure (Diehl et al., 2017). For each cell, spike times were circularly shifted by a single randomly selected temporal offset, thereby dissociating spike timing from animal position while preserving the temporal structure of the spike train. Spike times exceeding the session duration were wrapped to the beginning of the session. Spatial information was calculated for 1,000 shuffled datasets per cell. A neuron was classified as a potential place cell when its observed spatial information exceeded the 95^th^ percentile of its shuffled distribution. Place fields were further defined as contiguous regions ≥12 cm and ≤70 cm in length in which firing rate exceeded 20% of the peak firing rate. Cells with a peak firing rate <1 Hz were excluded from place field analysis. Recorded neurons were considered to be a place cell if they met the following criteria: i) information score ≥ 95^th^ percentile, ii) place field size ≥12 cm and ≤ 70 cm, and iii) peak firing rate > 1Hz. Within-session spatial stability was assessed by calculating the Pearson correlation coefficient between rate maps generated from odd and even numbered laps.

### Y-maze

Novelty discrimination was assessed using a three-arm Y-maze (Panlab, LE847) consisting of three identical enclosed arms (30 × 6 × 15 cm) arranged at 120° angles. Visual cues were positioned above and outside the maze. Room illumination was maintained at 20 lux. The maze orientation and entry arm were held constant across trials, while the identities of the familiar and novel arms were counterbalanced across animals. The test consisted of two trials separated by 60 min. During the exposure trial, mice were placed at the end of the entry arm and allowed to explore for 10 min with one arm inaccessible. Mice were then returned to their home-cage for 60 min. During the test trial, all three arms were accessible and mice were allowed to explore for 5 min. The maze was cleaned with 70% ethanol between trials. Behavior was recorded by video and tracked using DeepLabCut. Time spent in each arm was quantified using custom Python scripts. Novelty discrimination was quantified as a discrimination index (DI): DI = [*t*_novel_ – *t*_entry_]/[*t*_novel_ + *t*_entry_] × 100) or DI = [*t*_novel_ – *t*_familiar_]/[*t*_novel_ + *t*_familiar_] × 100) where *t* represents time spent in the indicated arm. Total distance traveled and mean running speed were calculated from smoothed x,y coordinates using a Gaussian filter (σ = 150 ms).

### Immunohistochemistry

Mice were transcardially perfused with 0.1 M PBS followed by 4% paraformaldehyde (PFA) in 0.1 M PBS, and brains were post-fixed overnight in 4% PFA. Coronal sections (50 μm, 300 µm apart) were prepared using a vibratome and processed using standard immunohistochemical procedures (Tierno et al., 2026). Primary antibodies included chicken anti-GFP (1:1,000; Aves, GFP1020; RRID: AB_10000240), rabbit anti-WFS1 (1:1,000; Proteintech, 11558-1-AP; RRID: AB_2216046), and mouse anti-GAD67 (1:1,000; Millipore, MAB5406; RRID: AB_2278725). Secondary antibodies were Alexa Fluor 488-conjugated goat anti-chicken IgG (A11039; RRID: AB_2534096), Alexa Fluor 546-conjugated goat anti-mouse IgG (A11030; RRID: AB_2737024), and Alexa Fluor 647-conjugated goat anti-rabbit IgG (A21244; RRID: AB_2535812), all at 1:1,000. Sections were mounted on charged slides with Fluoromount-G containing DAPI. Images were acquired using a Leica DM6 microscope at 5 to 20x with LAS X software and quantified using FIJI (ImageJ). Brightness and contrast were adjusted manually in FIJI (ImageJ) and applied uniformly across each image. For cell count quantification, all jGCaMP8m-expressing cells were quantified in three consecutive brain sections 300 µm apart through the injection site in CA1 per animal.

### Quantification and statistical analysis

Statistical analyses were performed using GraphPad Prism 11, Python 3.13.5 (SciPy 1.15.3), or MATLAB (MathWorks). Boxplots were generated using a seaborn boxplot function where boxes represent the interquartile range (IQR), the median indicated by a horizontal line; whiskers extend to the most extreme values within 1.5 × IQR, and individual points beyond this range are shown as outliers. Statistical tests included two-tailed Student’s *t*-tests, two-way repeated-measures ANOVA with Sidak or Tukey post hoc tests, Mann-Whitney *U* tests with Bonferroni correction, and Fisher’s exact test with Bonferroni correction. Data are presented as mean ± SEM unless otherwise indicated, and significance was set at *P* < 0.05.

## Results

### CA1 activity is preserved after brain injury

To determine whether chronic brain injury alters baseline CA1 network activity, we performed unilateral CCI over the somatosensory cortex of *Camk2a*-Cre mice at P60. Five weeks later, jGCaMP8m was expressed in dorsal CA1 of control and brain injured littermates, followed by implantation of GRIN lenses for one-photon *in vivo* calcium imaging (**Fig. 1A–C**). jGCaMP8m expression was largely restricted to WFS1-positive CA1 pyramidal neurons in both groups (**Fig. 1A,B**). Imaging was initiated 10-13 weeks after injury, and high quality neurons identified within the miniscope FOV were retained for analysis (**Fig. 1D,E**). The number of identified neurons was reduced in brain injured mice compared with controls (control: 192 ± 12; brain injury: 118 ± 11 cells; *P* = 4.87E-04, two-tailed *t*-test), possibly related to loss of CA1 neurons after brain injury (Saatman et al., 2006), whereas the proportion of identified cells meeting quality criteria for analysis was comparable between groups (control: 84 ± 2%; brain injury: 82 ± 3%; *P* = 0.48, Fisher’s exact test).

**Fig. 1.**
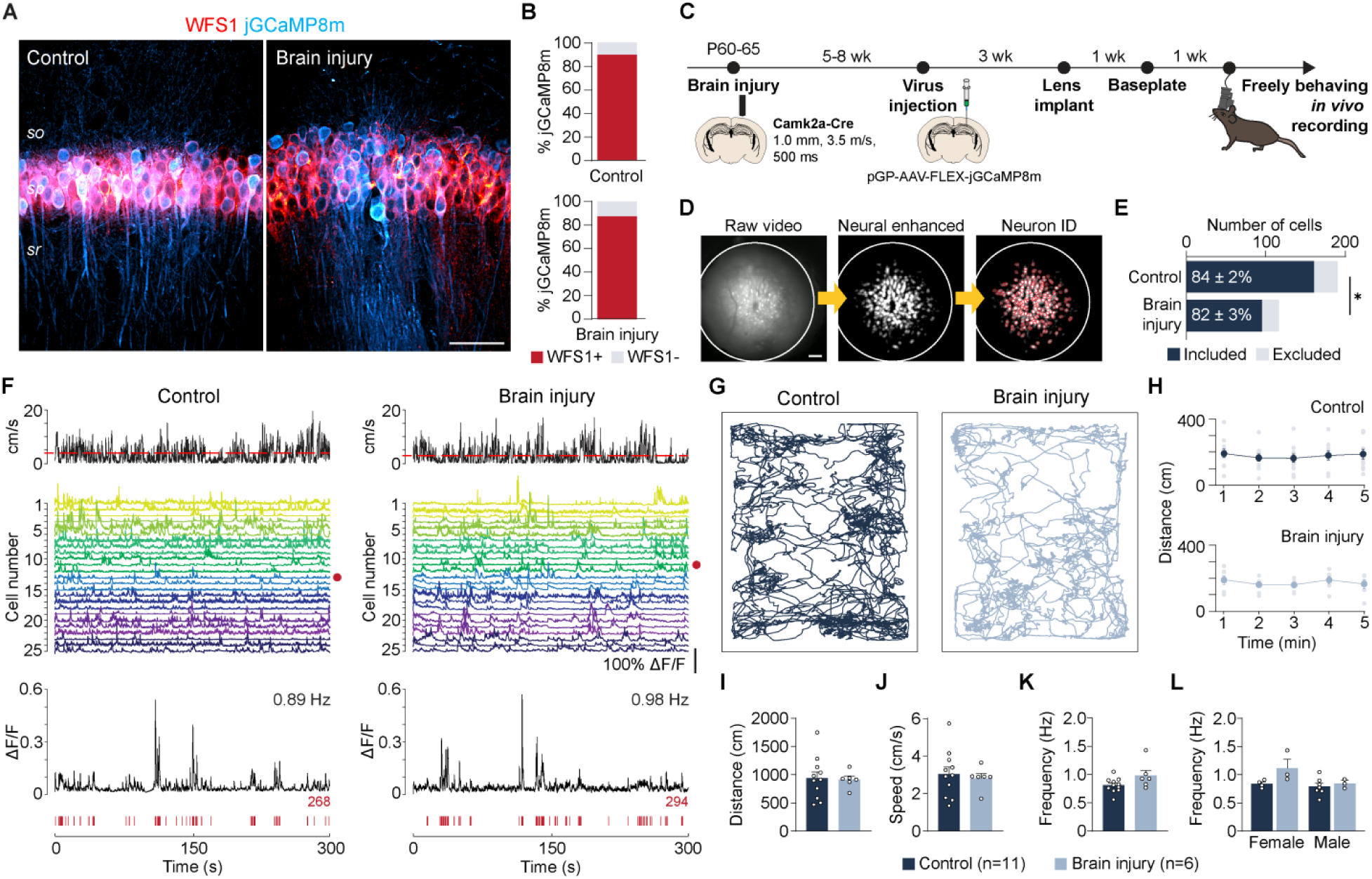
CA1 activity is preserved after brain injury. **(A)** Representative images of CA1 from control and brain injured *Camk2a*-Cre mice expressing FLEX-jGCaMP8m (blue) and immunolabeled for WFS1 (red). **(B)** Quantification of jGCaMP8m-expressing neurons co-labeled with WFS1 in control (top) and brain injured (bottom) mice. **(C)** Experimental timeline for *in vivo* calcium imaging. **(D)** Representative steps in calcium signal processing using MIN1PIPE. Left, raw miniscope FOV; middle, neural enhanced image; right, identified neuronal contours (red). **(E)** Number of neurons identified in the miniscope FOV in control and brain injured mice (control, 192 ± 12 cells, *n* = 11 mice; brain injury, 118 ± 11 cells, *n* = 6 mice; *P* = 4.87E-04, two-tailed *t*-test). The percentage of identified neurons included in subsequent analyses is indicated within each bar; exclusion criteria are described in Figure S3. **(F)** Representative home-cage recordings from control and brain injured mice. Top, locomotor speed; red dashed line indicates mean speed during the recording session. Middle, calcium traces from 25 representative neurons. Bottom, representative calcium trace and inferred spike events (red) from the neuron indicated by the red marker above. **(G)** Representative home-cage trajectories from control and brain injured mice. **(H)** Distance traveled over consecutive 1 min intervals during the recording session. Lines and symbols indicate group means; individual mouse values are shown in gray. **(I)** Total distance traveled during the recording session. **(J)** Mean locomotor speed. **(K)** Mean CA1 calcium event frequency per mouse. **(L)** Mean calcium event frequency separated by sex. Data are mean ± SEM. Scale bars, 50 μm (A) and 100 μm (D). SO, stratum oriens; SP, stratum pyramidale; SR, stratum radiatum.

We then quantified CA1 neuronal activity during home-cage exploration using spike inference from calcium traces (**Fig. 1F**). Calcium event frequency did not differ between control and brain injured mice, either overall or when analyzed by sex (**Fig. 1K,L**). Home-cage behavior was also comparable between groups, including movement trajectories, total distance traveled and average speed (**Fig. 1G–J**). Thus, despite a reduction in the number of neurons detected within the imaging FOV after injury, baseline CA1 neuron activity and locomotor behavior were broadly preserved during the chronic phase of TBI.

### Brain injury reduces the precision of CA1 spatial activity

Next, we asked whether chronic brain injury alters the organization of spatially responsive CA1 neurons. Mice were trained to run back and forth on a 2 m linear track for 7 days, with four sessions per day (**Fig. 2A**). By day 7, both groups were able to complete at least 15 laps within a 10 min session (**Fig. 2B,C**). Running speed did not differ between control and brain injured mice across sessions, although both groups consistently ran faster in the leftward direction (**Fig. S2**). Calcium traces were extracted and aligned to the animal’s position and running direction. Spatially tuned neurons were identified in both groups and during both running directions, including neurons with directionally selective or bidirectional place fields as well as neurons without spatially tuned activity (**Fig. 2D–G; Fig. S5**). Because place cell activity is directionally organized in one-dimensional environments (Muller et al., 1994), rightward and leftward running were analyzed independently. Place cells were identified using a single-cell, shuffle-based spatial information criterion (Skaggs et al., 1993; Diehl et al., 2017), with neurons classified as place cells when their spatial information exceeded the 95^th^ percentile of their shuffled distribution (**Fig. 2H**), field size was 12 – 70 cm and peak firing rate > 1Hz.

**Fig. 2.**
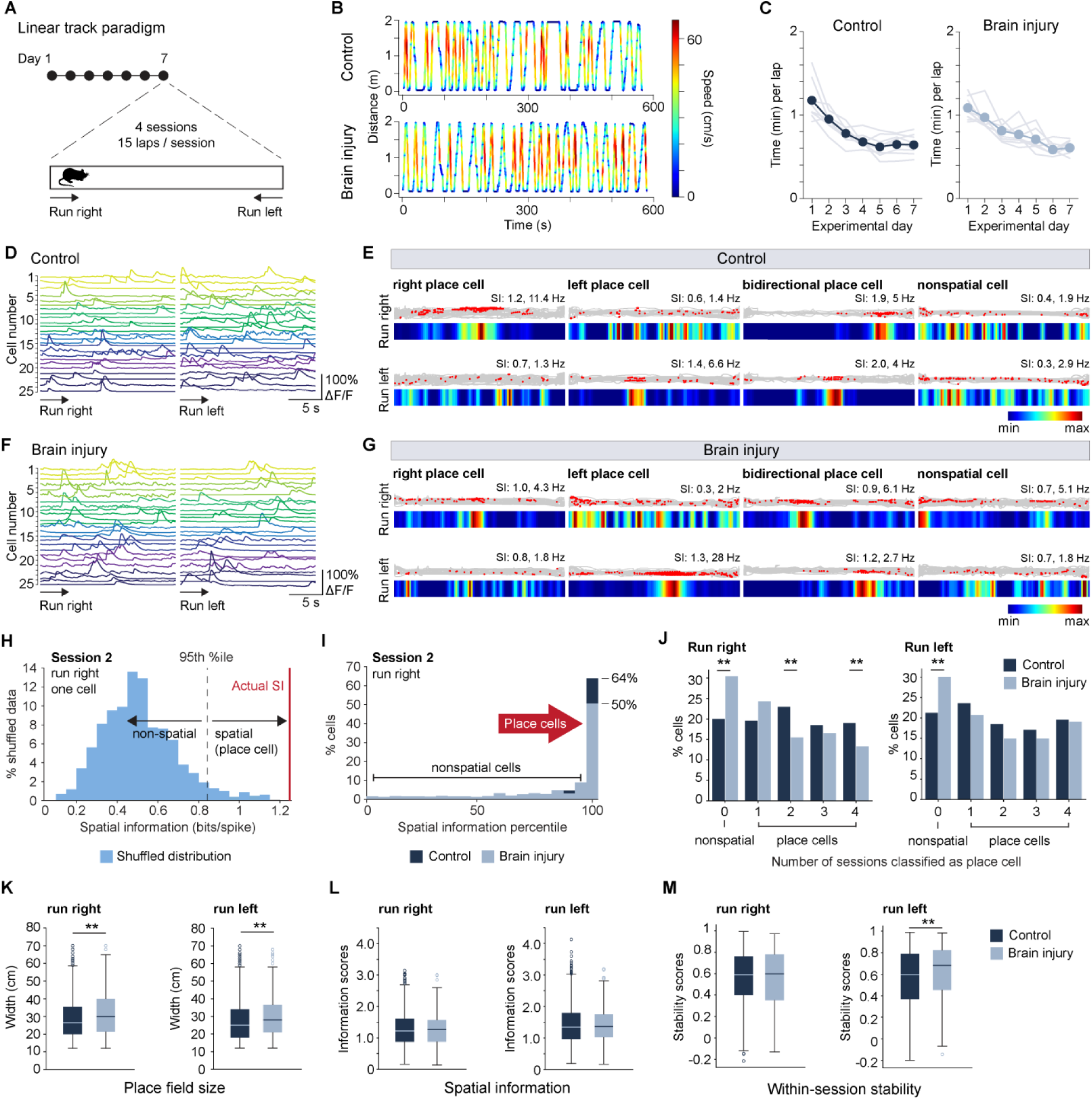
Chronic brain injury alters the recruitment and spatial precision of CA1 representations. **(A)** Experimental design for the linear track task. **(B)** Representative running trajectories and speed profiles from control and brain injured mice. **(C)** Time per lap across experimental days. Gray lines indicate individual mice and colored symbols indicate group means. **(D, F)** Representative calcium traces during rightward and leftward running from control (D) and brain injured (F) mice. **(E, G)** Representative rightward, leftward, bidirectional, and nonspatial cells from control (E) and brain injured (G) mice. Gray trajectories indicate position and red dots indicate inferred calcium events; spatial firing rate maps, spatial information (SI), and peak event frequency are shown for each running direction. **(H)** Example shuffled SI distribution used for place cell classification. **(I)** Distribution of SI percentiles for all cells during one session; the 95^th^ percentile threshold and proportion of place cells are indicated. **(J)** Percentage of cells classified as place cells according to the number of sessions in which they met place cell criteria. Run right: *P* = 1.11E-06, 3.2E-04, and 6.27E-03 for 0, 2 and 4 sessions; run left: *P* = 6.9E-05 for 0 sessions; Fisher’s exact test with Bonferroni correction. **(K)** Place field width. Run right, *P* = 2.72E-04; run left, *P* = 4.29E-05. **(L)** Mean SI per group. **(M)** Within-session stability. Run left, *P* = 2.28E-05. Mann–Whitney U test for K–M. *n* = 1,205 cells from 8 control mice and 455 cells from 6 brain injured mice for K–M; *n* = 1,506 cells from 8 control mice and 654 cells from 6 brain injured mice for J. Box plots show median and interquartile range (IQR), with whiskers extending to 1.5 × IQR.

Although brain injured mice retained neurons that met place cell criteria, a significantly greater proportion of neurons never qualified as a place cell across the four imaging sessions, while fewer neurons were repeatedly classified as place cells (**Fig. 2I,J**). This reduction in place cell recruitment was evident during both running directions. Thus, chronic brain injury reduces the population of CA1 neurons that participates in spatial representations across repeated experience.

We next examined the properties of neurons that did form place fields. Place fields were modestly but consistently broader in brain injured mice during both rightward and leftward running (**Fig. 2K**). In contrast, spatial information was preserved among place cells and did not differ between groups (**Fig. 2L**). Within-session stability was also preserved during rightward running and was modestly increased in brain injured mice during leftward running (**Fig. 2M**). Stability increased with repeated exposure to the track in both groups and running directions (**Fig. S6A,B**), and peak and mean firing rates of place cells were comparable between groups (**Fig. S6C,D**). Thus, brain injury does not abolish the spatial activity of established place cells. Rather, neurons that form place fields exhibit broader spatial tuning despite preserved spatial information and within-session stability.

Nonspatial neurons in brain injured mice had lower firing rates than controls during both running directions, and their within-session stability was also reduced (**Fig. S6E-H**). This suggests neurons that failed to meet place cell criteria are not simply inactive; they exhibited reduced and less stable activity after injury. Together with the reduced recruitment of neurons into place cell populations, our findings indicate that chronic brain injury alters the cellular organization of hippocampal spatial activity, with fewer neurons participating in spatial representations, broader spatial tuning among recruited neurons, and reduced activity and stability among neurons that remain nonspatial.

### Brain injury impairs spatial novelty discrimination

To determine whether altered organization of CA1 spatial activity was accompanied by impaired spatial memory, we tested the same mice in a Y-maze novelty discrimination task (**Fig. 3A**). Mice were initially restricted to two arms of the maze and, following a 1 h rest period in the home-cage, were allowed to explore all three arms. During the test phase, control mice preferentially explored the novel arm, spending more time there than brain injured mice, which showed no preference among the three arms (**Fig. S7A**). To quantify novelty discrimination, we calculated discrimination indices comparing time spent in the novel arm with time spent in either the entry or familiar arm (**Fig. 3B**). Control mice exhibited significantly greater discrimination indices for both comparisons than brain injured mice, consistent with increased preference for the novel arm. These differences were not attributable to altered locomotor behavior, as total distance traveled and running speed were comparable between groups (**Fig. S7B,C**). Thus, chronic brain injury impairs discrimination of a novel spatial environment despite preserved locomotor performance.

**Fig. 3.**
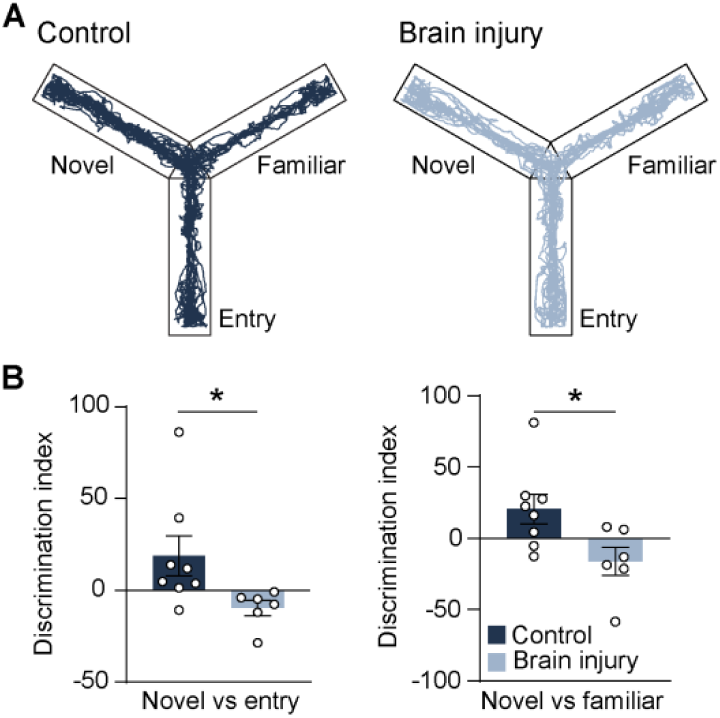
Novelty discrimination is impaired after brain injury. **(A)** Representative tracking plots from control (left) and brain injured (right) mice during the test phase of the Y-maze task. **(B)** Discrimination index for the novel arm relative to the entry arm (left) or familiar arm (right) (novel vs. entry: control, 18.8 ± 10.9; brain injury, −9.7 ± 4.1; *P* = 0.04; novel vs. familiar: control, 20.5 ± 10.2; brain injury, −16.1 ± 9.8; *P* = 0.02; two-tailed *t*-test, *n* = 8 control and 6 brain injured mice). Data are mean ± SEM.

## Discussion

TBI produces substantial remodeling of hippocampal excitatory and inhibitory networks, with changes predicted to alter CA1 excitability in opposing directions. Loss of inhibition and the emergence of aberrant recurrent glutamatergic circuits could promote elevated excitability (Toth et al., 1997; Santhakumar et al., 2000; Hunt et al., 2009, 2011), whereas overall synapse loss and reductions in long-range afferent inputs could constrain neuronal activity (Scheff et al., 2005; Frankowski et al., 2022). Despite these competing influences, we found CA1 firing was broadly preserved three months after injury, a finding that has been consistently reported in models of TBI (Eakin and Miller, 2012; Koch et al., 2020; Ulyanova et al., 2023; Adam et al., 2026). Thus, post-traumatic circuit remodeling does not produce a simple change in the overall level of CA1 activity or excitation–inhibition balance, but rather alters how that activity is organized across neuronal populations. The spatial representation of the environment provides a particularly clear example of this reorganization. TBI increases the fraction of neurons that remain spatially unresponsive, broadens the place fields of neurons that do form spatial representations and alters the activity and stability of the nonspatial population.

Despite these changes, several properties of established place cell representations remain intact. Spatial information and within-session stability were preserved among neurons that formed place fields, indicating that chronic TBI does not abolish the capacity of CA1 neurons to encode space. This may represent a degree of functional compensation that preserves fundamental hippocampal processing without restoring the fidelity required for normal memory. Several features of post-traumatic circuit remodeling could contribute to this incomplete compensation. Local inhibition constrains the formation and spatial tuning of CA1 place fields (Royer et al., 2012; Dudok et al., 2021; Geiller et al., 2022) and is prominently disrupted after TBI (Hunt et al., 2011; Ulyanova et al., 2023), potentially broadening spatial tuning and impairing the recruitment of neurons into spatial representations. TBI also disrupts long-range inputs from entorhinal cortex (Frankowski et al., 2022), which could reduce the spatial information available to CA1. Consistent with this possibility, disruption of medial entorhinal input produces broader and less precise CA1 place fields (Hales et al., 2014). The reduced firing and stability of nonspatial neurons further suggest that altered inhibitory or afferent inputs may affect both the formation of spatial representations and the precision of established place fields. Intriguingly, transplantation of inhibitory neurons into the hippocampus improves memory after TBI (Zhu et al., 2019), while transplanted neurons also receive new entorhinal inputs (Frankowski et al., 2022), providing evidence that restoration of local inhibition and afferent connectivity may improve hippocampal function after TBI. At the same time, post-traumatic circuit remodeling may carry pathological consequences, as loss of inhibition, enhanced excitatory connectivity and aberrant recurrent circuit formation can promote network hyperexcitability and post-traumatic epilepsy (Hunt et al., 2009). Although epilepsy was not assessed in this study, no seizures were observed during experimentation.

The spatial phenotype we observed after chronic TBI also differs from epilepsy and Alzheimer’s disease models. In epilepsy, place cell representations show reductions in the number of spatially tuned neurons, spatial information and stability, with pronounced disruption of the temporal organization of place cell firing (Liu et al., 2003; Shuman et al., 2020). Alzheimer’s disease models similarly exhibit reduced recruitment and spatial information without a change in field size (Mably et al., 2017; Jun et al., 2020). Thus, whereas these models show a broader degradation of established spatial representations, chronic TBI produces a more selective reorganization of the cellular composition and spatial tuning of CA1 representations. These differences may reflect the more restricted spatial extent of CCI pathology compared with the widespread CNS pathology produced by status epilepticus or the distributed cortical–hippocampal pathology of Alzheimer’s disease models. Although CCI can induce secondary circuit remodeling beyond the primary lesion (Holden et al., 2019; Frankowski et al., 2022), the relative preservation of spatial representations may reflect this more focal pattern of injury. Thus, chronic post-traumatic remodeling may preserve fundamental aspects of hippocampal network function while compromising the fidelity with which they are implemented.

Together, our findings support a model in which TBI preserves overall CA1 activity while reorganizing the cellular and spatial structure of hippocampal representations. This dissociation between neuronal activity and network function provides a circuit-level framework for understanding how persistent post-traumatic remodeling can preserve basic hippocampal activity while compromising the precision required for spatial memory.

## Author contributions

A.T. contributed to design, execution and analysis of experiments, funding and wrote the manuscript. R.F.H. contributed to the concept, design, analysis of experiments, funding and edited the manuscript.

## Acknowledgments

We thank Di Wu for manual inspection of automated spike inferences and Daniel Aharoni for recommendations on analysis pipelines. This work was supported by funding from the National Institutes of Health grants F31–NS132447 (A.T.) and R01–NS096012 (R.F.H.).

## Supplementary figures

**Fig. S1.**
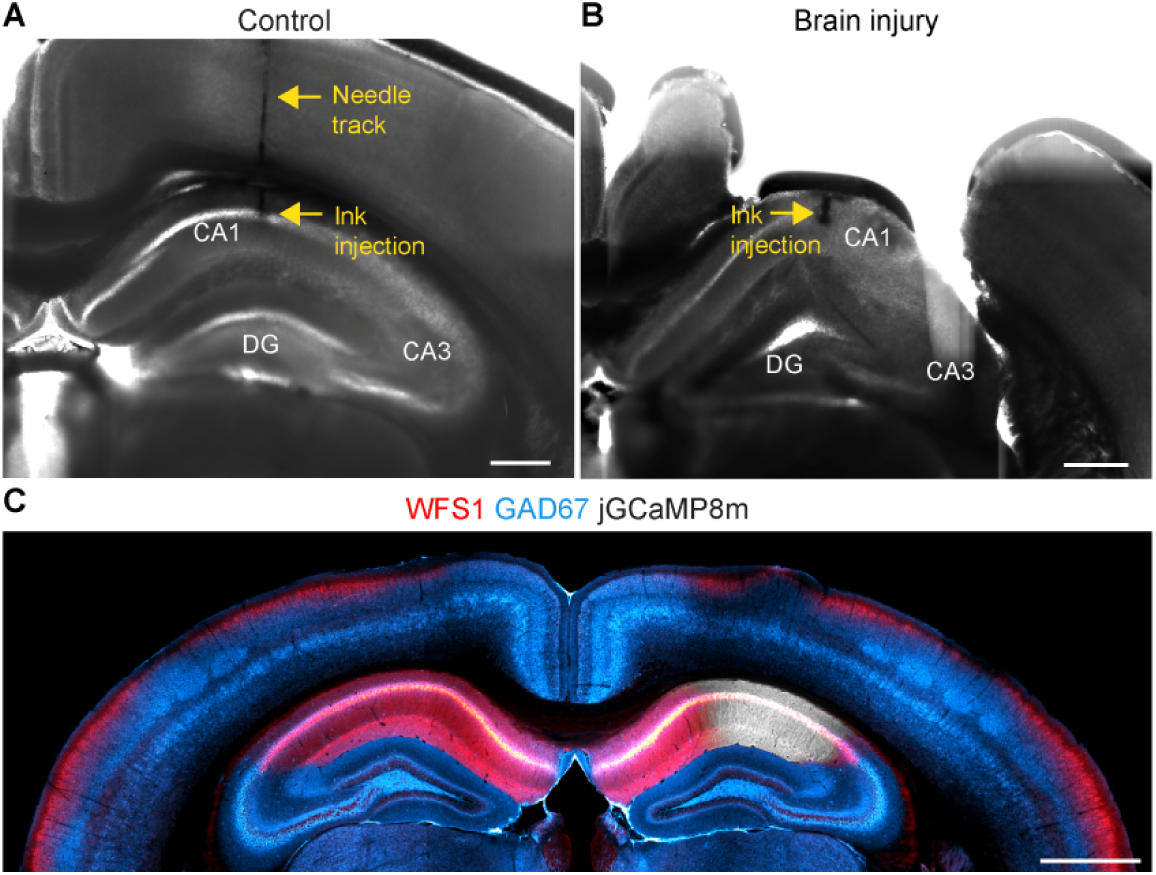
Verification of CA1 targeting and cell type specificity of jGCaMP8m expression. **(A)** Representative coronal section from a control mouse following ink injection into the CA1 region. Yellow arrows indicate the needle track and injection site. **(B)** Representative coronal section from a brain injured mouse following ink injection into the CA1 region. Yellow arrow indicates the injection site. **(C)** Representative coronal section from a control *Camk2a*-Cre mouse expressing jGCaMP8m (white) in CA1 and co-labeled for WFS1 (red) and GAD67 (blue). Scale bars, 500 μm (A, B) and 1 mm (C).

**Fig. S2.**
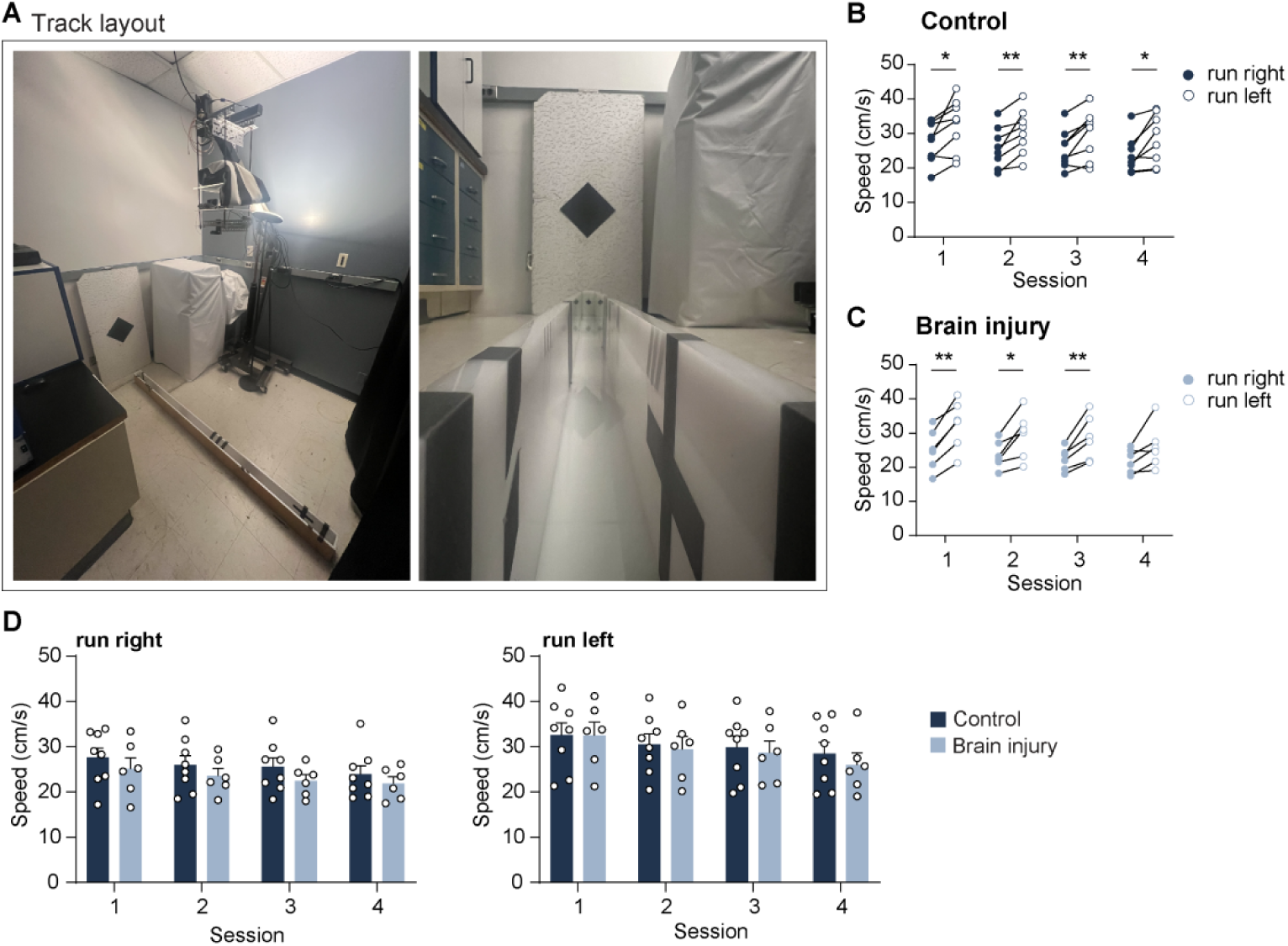
Track layout and running speed. **(A)** Linear track apparatus and environmental cues. Mice ran between the two ends of the track, with running direction defined as rightward or leftward. **(B, C)** Speed (cm/s) during rightward (closed circles) and leftward (open circles) running across four sessions in control (B) and brain injured (C) mice. *P* values indicate rightward versus leftward running within each session (control: session 1, *P* = 0.02; session 2, *P* = 5E-03; session 3, *P* = 8E-03; session 4, *P* = 0.02; brain injury: session 1, *P* = 1E-03; session 2, *P* = 0.02; session 3, *P* = 8E-03; session 4, *P* = 0.06; two-way repeated measures ANOVA with Tukey’s post hoc test; *n* = 8 control and 6 brain injured mice). **(D)** Mean running speed across sessions for control and brain injured mice during rightward (left) and leftward (right) running. Data points represent individual mice. Error bars, SEM.

**Fig. S3.**
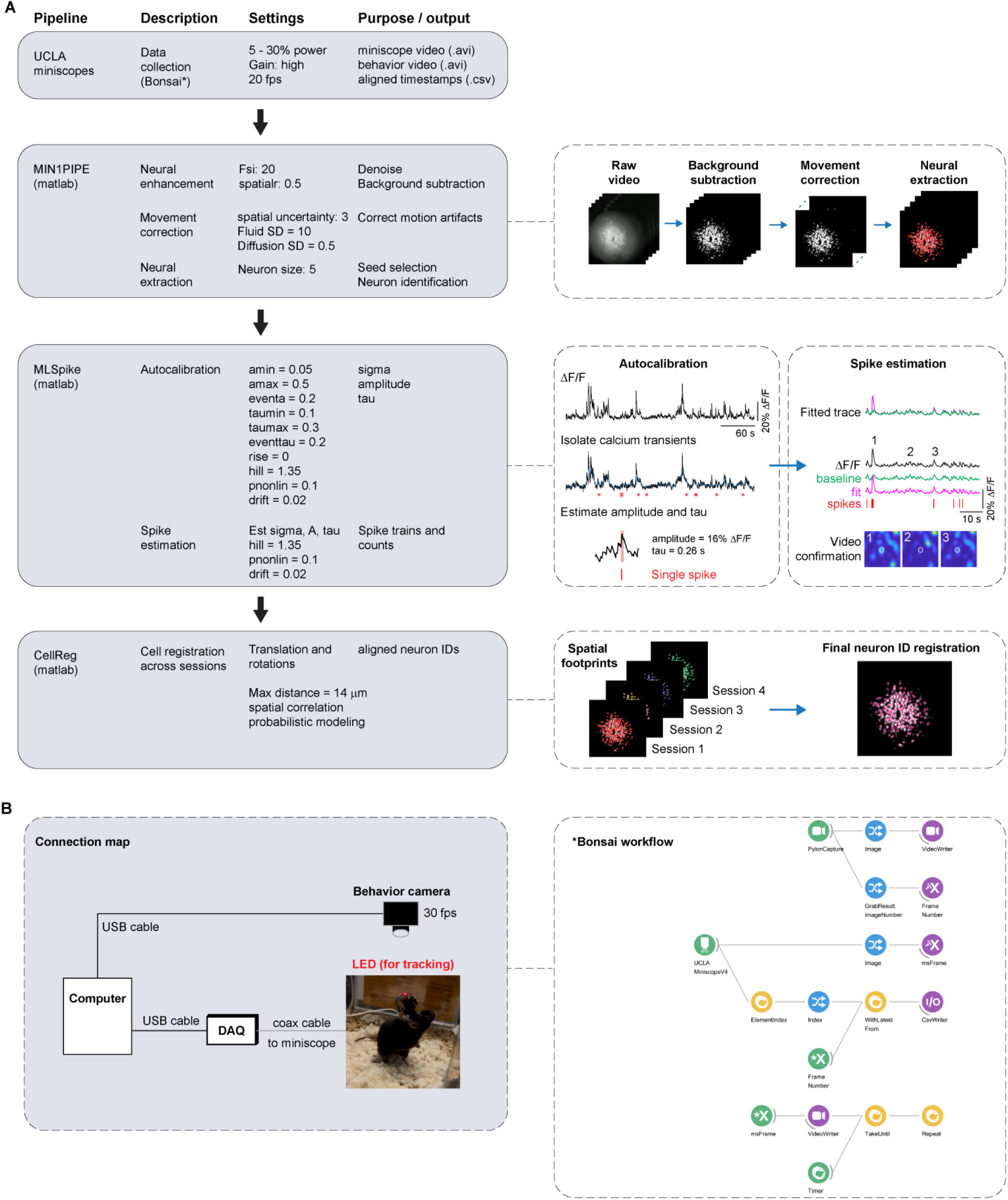
Pipeline for calcium signal extraction and analysis from miniscope recordings. **(A)** Overview of the calcium imaging and analysis pipeline. Miniscope recordings were acquired using a UCLA Miniscope and synchronized with behavioral video using a custom Bonsai workflow. Calcium imaging videos were processed using MIN1PIPE for background subtraction, movement correction, neural extraction, and neuron identification. Calcium traces were then processed using MLspike for spike estimation, including autocalibration of single spike amplitude and decay parameters followed by estimation of spike events across the full trace. Trace alignment was manually verified and corrected for each neuron and mouse. For longitudinal linear track recordings, neurons were registered across sessions using CellReg, and registered neuron identities were used for downstream analyses. **(B)** Miniscope recording and synchronization setup. Left, connections between the computer, behavior camera, data acquisition device, and miniscope. Right, Bonsai workflow used for real-time synchronization of behavioral and calcium imaging recordings. Outputs include miniscope video (.avi), behavioral video (.avi), and aligned frame timestamps (.csv).

**Fig. S4.**
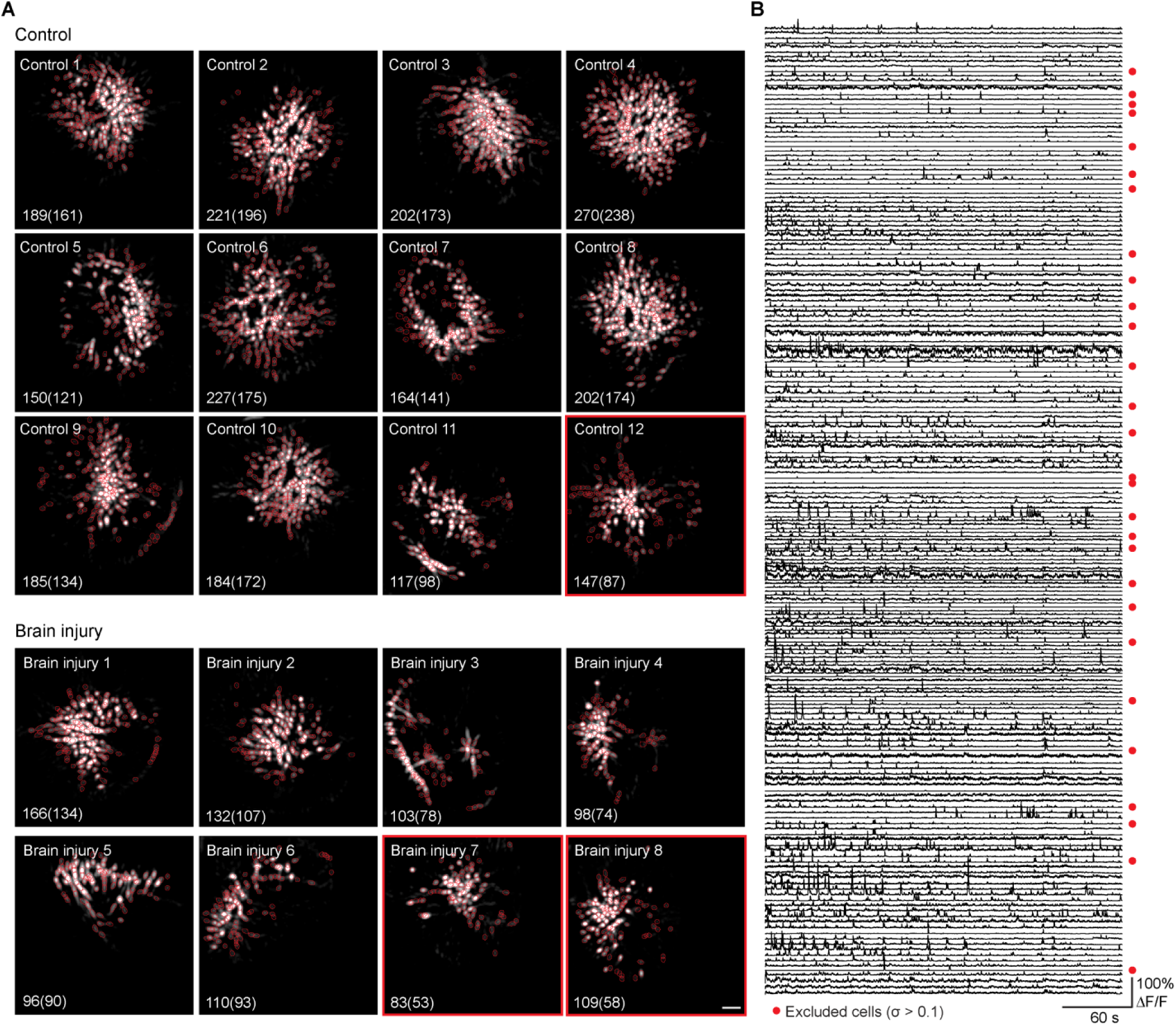
Field of view (FOV) and calcium traces from all recorded mice. **(A)** Miniscope FOVs from all recorded mice following calcium signal extraction with MIN1PIPE. Individual neuronal contours are outlined in red. Red boxes indicate mice excluded from analysis because >30% of extracted neurons had high noise (σ > 0.1). Numbers indicate the total number of identified neurons, with the number included in subsequent analyses shown in parentheses. **(B)** Calcium traces from all neurons identified in one representative control mouse during the home-cage recording. Red circles indicate neurons excluded from analysis based on the noise criterion. Scale bar, 100 μm.

**Fig. S5.**
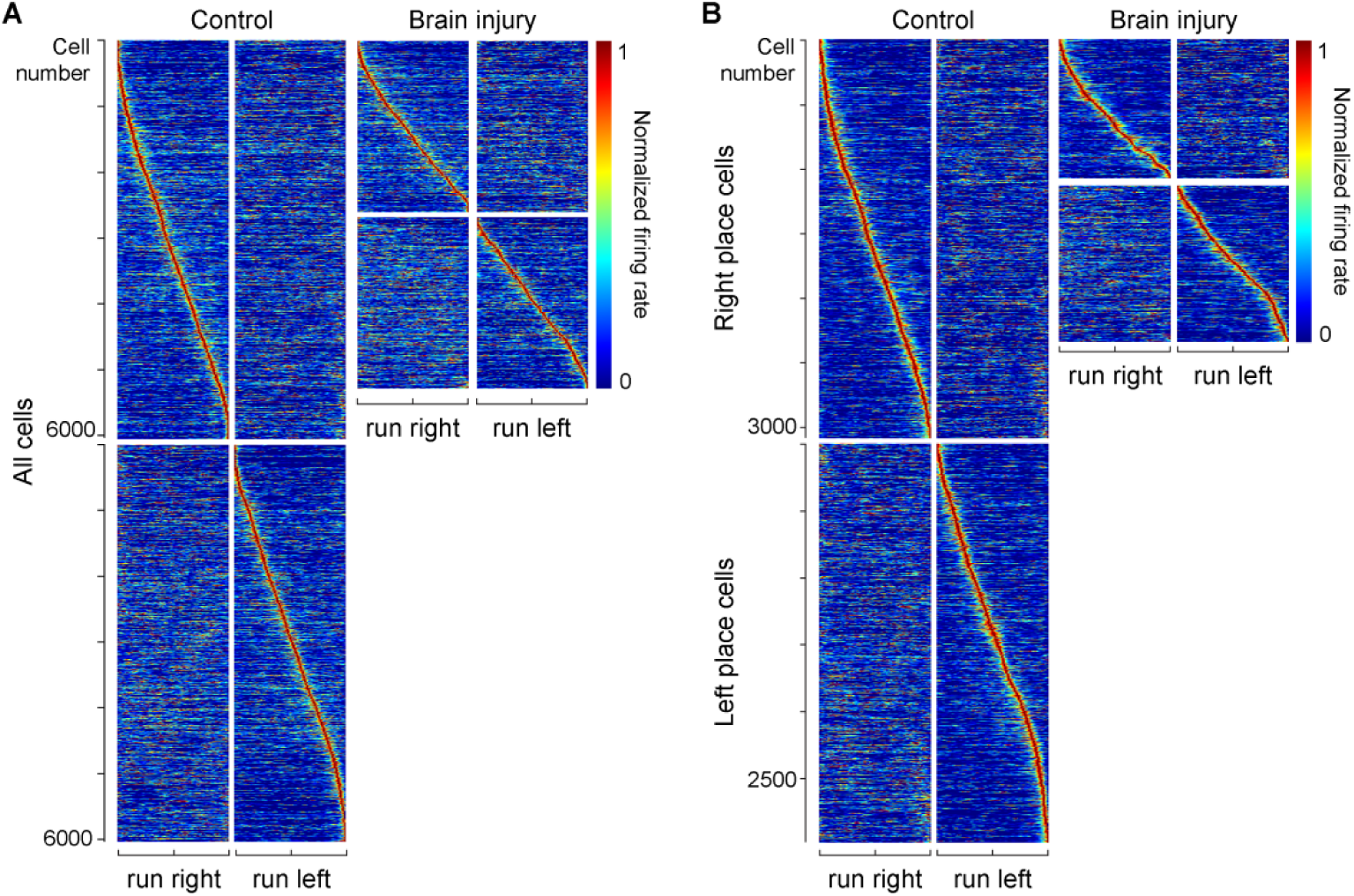
Firing rate maps of CA1 neurons. **(A)** Spatial firing rate maps for all cells recorded across four linear track sessions during rightward and leftward running in control and brain injured mice. Maps are ordered by peak firing rate for each running direction (*n* = 6,024 cells from 8 control mice; *n* = 2,616 cells from 6 brain injured mice). **(B)** Spatial firing rate maps for place cells recorded across four sessions during rightward and leftward running. Maps are ordered by peak firing rate (*n* = 3,078 rightward and 2,972 leftward place cells from 8 control mice; *n* = 1,074 rightward and 1,166 leftward place cells from 6 brain injured mice). Firing rates are normalized to the maximum firing rate of each cell.

**Fig. S6.**
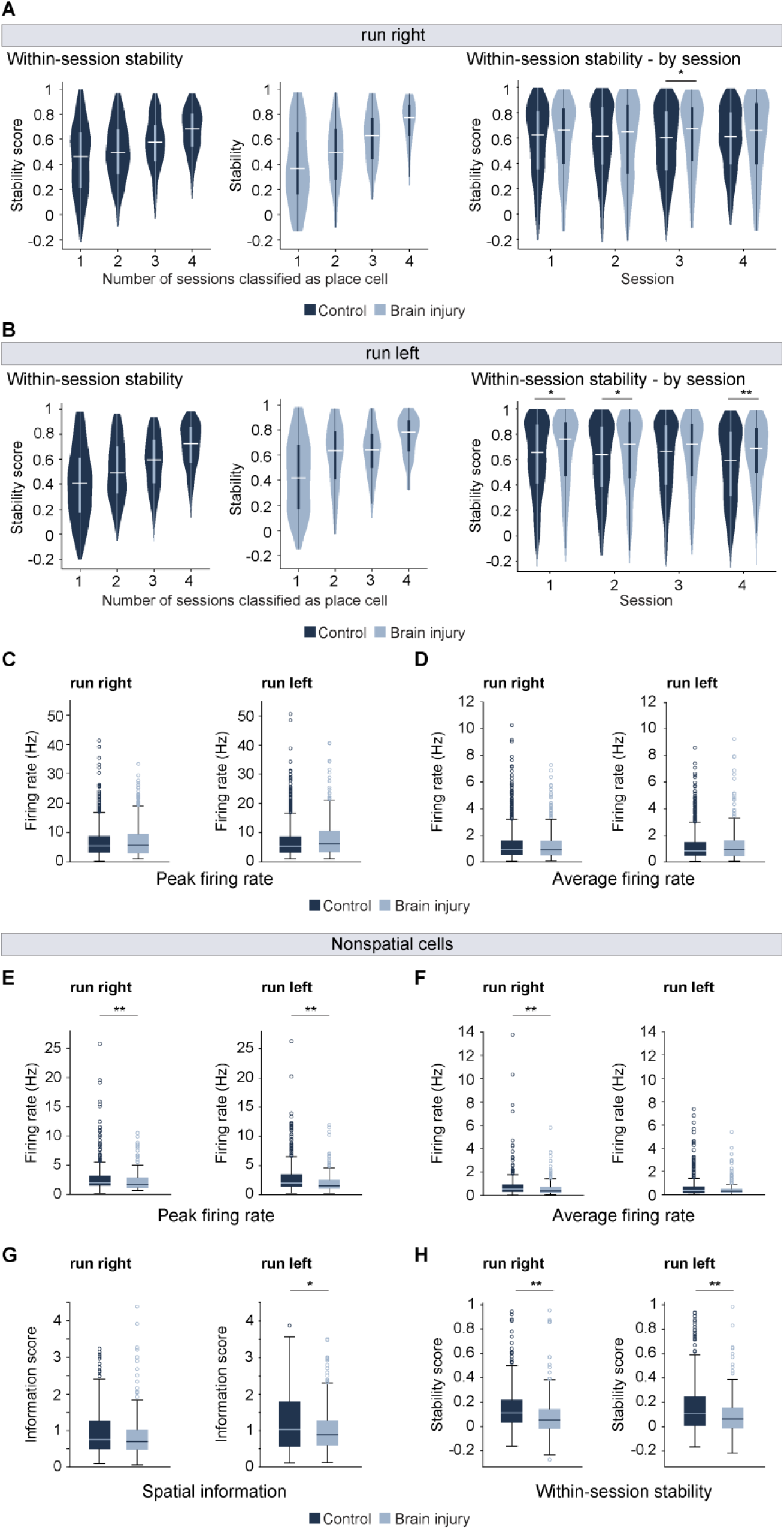
Place cell stability, firing rates and nonspatial cell properties. **(A)** Within-session stability of place cells during rightward running as a function of the number of sessions in which cells were classified as place cells (left) and by session number (right) in control and brain injured mice (session 3, *P* = 0.02; Mann–Whitney *U* test with Bonferroni correction). **(B)** Same as **A** for leftward running (session 1, *P* = 0.04; session 2, *P* = 0.016; session 4, *P* = 7.3E-05; Mann–Whitney *U* test with Bonferroni correction). For A, B, *n* = 3,078 rightward place cells from 8 control mice and 1,074 from 6 brain injured mice; *n* = 2,972 leftward place cells from 8 control mice and 1,166 from 6 brain injured mice. Violin plots show the distribution of stability values; white lines indicate medians and boxes indicate interquartile ranges (IQR). **(C)** Peak firing rates of place cells during rightward and leftward running. **(D)** Mean firing rates of place cells during rightward and leftward running. For C, D, *n* = 1,205 cells from 8 control mice and 455 cells from 6 brain injured mice. **(E)** Spatial information of nonspatial cells, defined as cells that were not classified as place cells in any session, during rightward and leftward running (leftward, *P* = 1.55E-02). **(F)** Within-session stability of nonspatial cells (rightward, *P* = 5.52E-06; leftward, *P* = 6.41E-04). **(G)** Peak firing rate of nonspatial cells (rightward, *P* = 4.45E-04; leftward, *P* = 3.36E-07). **(H)** Mean firing rate of nonspatial cells (rightward, *P* = 1.82E-04). Mann–Whitney *U* test was used for E–H. For nonspatial cells, *n* = 301 control and 199 brain injured cells for rightward running and 320 control and 197 brain injured cells for leftward running. Box plots show median, IQR, and whiskers extending to 1.5 × IQR.

**Fig. S7.**
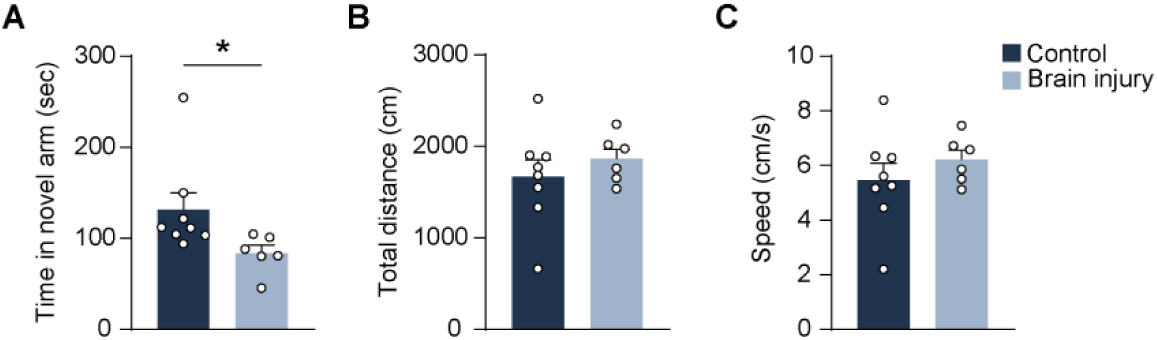
Performance during the Y-maze task. **(A)** Time spent in the novel arm during the test phase in control and brain injured mice (control, 131.4 ± 18.6 s; Brain injury, 83.4 ± 8.6 s; \**P* = 0.04, two-tailed *t*-test). **(B)** Total distance traveled during the test phase in control and brain injured mice (control, 1663 ± 187.9 cm; brain injury, 1862 ± 106.7 cm). **(C)** Mean locomotor speed during the test phase (control, 5.5 ± 0.6 cm/s; brain injury, 6.2 ± 0.4 cm/s). Data are mean ± SEM.

